# A gut commensal protozoa remotely shapes a lung niche for asthma-promoting eosinophils

**DOI:** 10.1101/2022.03.25.485893

**Authors:** Kyle Burrows, Louis Ngai, Pailin Chiaranunt, Eric Cao, Catherine J. Streutker, Brian Forde, Liam O’Mahony, Arthur Mortha

## Abstract

The gut microbiome influences chronic inflammation of the airways via the gut-lung axis. However, causal connections between microbes and their host, including the underlying mechanisms for this phenomenon remain largely unknown. Here, we show that colonization with the gut commensal protozoa, *Tritrichomonas musculis* (*T.mu*), remotely shapes the lung immune landscape and exacerbates allergic airway inflammation. We demonstrate that colonization with *T.*mu mediates the T and B cell-dependent accumulation and activation of inflammatory group 2 innate lymphoid cells in the lungs to constitute a tripartite immune network that serves as a niche for lung eosinophils. Animals colonized with *T.mu* show severely exacerbated allergic inflammation in the airways and reveal a new protozoan-driven gut-lung axis that remotely shapes the lung immune network to potentiate chronic pulmonary inflammation.

**One-Sentence Summary:** A gut microbe exacerbates asthma severity by promoting lung eosinophilia through a tripartite lymphocyte immune network.

---

The intestinal microbiota, the collection of microbes living in our gut, has profound impact on extra-intestinal organ homeostasis and inflammation (*1*). Asthma, a heterogeneous chronic inflammatory condition of the airways and lungs, is characterized by substantial tissue remodeling and inflammation (*2*). Increasing experimental and clinical evidence suggest that the gut microbiota serves as an associated risk factor for the development and severity of asthma (*3*). However, current data merely implicates bacterial dysbiosis during asthma, lacking the identification and proof of causally associated microbial species with the risk for developing asthma (*4*). Unlike pathogens that can directly infect the lung, commensal members of the gut microbiota are suspected to elicit their extra-intestinal influences across gut-tissue axes (*5*). The underlying mechanisms determining tissue specificity of this microbial-driven axes remain unclear and may involve microbial metabolites or the modulation of neuronal innervation (*6, 7*). Migration of immune cells further poses as attractive mechanism by which the microbiome could promote extra-intestinal immune manifestations. Group 2 innate lymphoid cells (ILC2), implicated in the development asthma, were shown to egress the intestine and transiently accumulate in the lung during helminth infections (*8*).

Natural (n)ILC2s are tissue-resident cells of mucosal organs including the intestine and lung, characterized by the lack of lineage markers, the expression of GATA3 and production of interleukin (IL)-5 and IL-13. Multiple signals, including the tuft-cell derived cytokine IL-25, can activate ILC2 (*9*). Anti-helminth immunity relies on IL-25 dependent ILC2 activation to expand mucus-releasing goblet cells, the accumulation of eosinophils and mechanical expulsion of the parasites through smooth muscle contractions (*10*). Stimulation of taste receptors on tuft cells promotes the release of IL-25 and drives the local expansion of gut-resident, inflammatory (i)ILC2 that egress from the intestine into the lymphatic system, resulting in the temporary accumulation of these highly activated innate immune cells in the lung (*8, 11*).

Others and our group identified *Tritrichomonas* spp. (*T.mu, T.muris* and *T.rainier*) as protozoan commensals of the murine intestinal tract (*12–14*). *Pentatrichomonas hominis* and *Dientamoeba fragilis* are two human-associated, close relatives of murine *Tritrichomonas* spp. with a restricted preference to colonize the intestine (*12*). Pertinently, both human and mouse protozoa show signs of commensalism due to the lack of clinical manifestation across multiple colonized, yet “healthy” human cohorts and mouse line, suggest that murine *Tritrichomonas* spp. may be considered commensals rather than pathobionts (*15*). Improved anti-helminth and anti-bacterial immunity following engraftment of *Tritrichomonas* spp. require the activation of the immune system through ATP-dependent NLRP1B and NLRP3 activation or succinate-mediated release of IL-25 by tuft cells (*12–14, 16*).

Colonization with *Tritrichomonas* spp. promotes a tuft cell-driven activation of ILC2 in the gut and leads to the local accumulation of eosinophils (*14*). Both ILC2 and eosinophils are abundantly found in the lungs of mice with allergic asthma, raising the hypothesis that commensal *Tritrichomonas* spp. promote a remote regulation of extra-intestinal organs like the lung (*17*). Due to its previously demonstrated ability to drive ILC2 activation and goblet cell hyperplasia in an IL-25-dependent fashion, we examined whether the protozoan commensal *Tritrichomonas musculis* (*T.mu*) can serve as remote regulator of type 2 immune cell activation at extra-intestinal sites across gut-tissue axis. Here, we show that the gut commensal protist *T.mu* remotely acts as regulator of a lung niche for asthma-promoting eosinophils extending our knowledge of how the gut microbiota controls extra-intestinal tissue immunity and remote-regulated disease outcomes.

To investigate whether colonization with *T.mu* affects type 2 immunity beyond the intestine, multiple extra-intestinal tissues from colonized mice were analyzed for eosinophilia. We observed increased frequencies of eosinophils in the blood, colon, bone marrow and lung three weeks post colonization by *T.mu* (Fig.1A-C and fig 1a), followed by a concomitant elevation of IL-5 receptor (CD125)-expressing eosinophil precursors in the bone marrow (BM)(fig.1b). Colonization by the protozoa selectively affected eosinophilopoiesis and did not alter the numbers of hematopoietic stem cells (LSK; lineage(Lin)^-^Sca-1^+^c-Kit^+^) or other myeloid precursor populations (i.e., granulocyte macrophage precursors (GMP)) (fig.1b), implicating a systemic increase in eosinophils and eosinophilopoiesis after colonization with *T.mu*.

**Figure 1.**
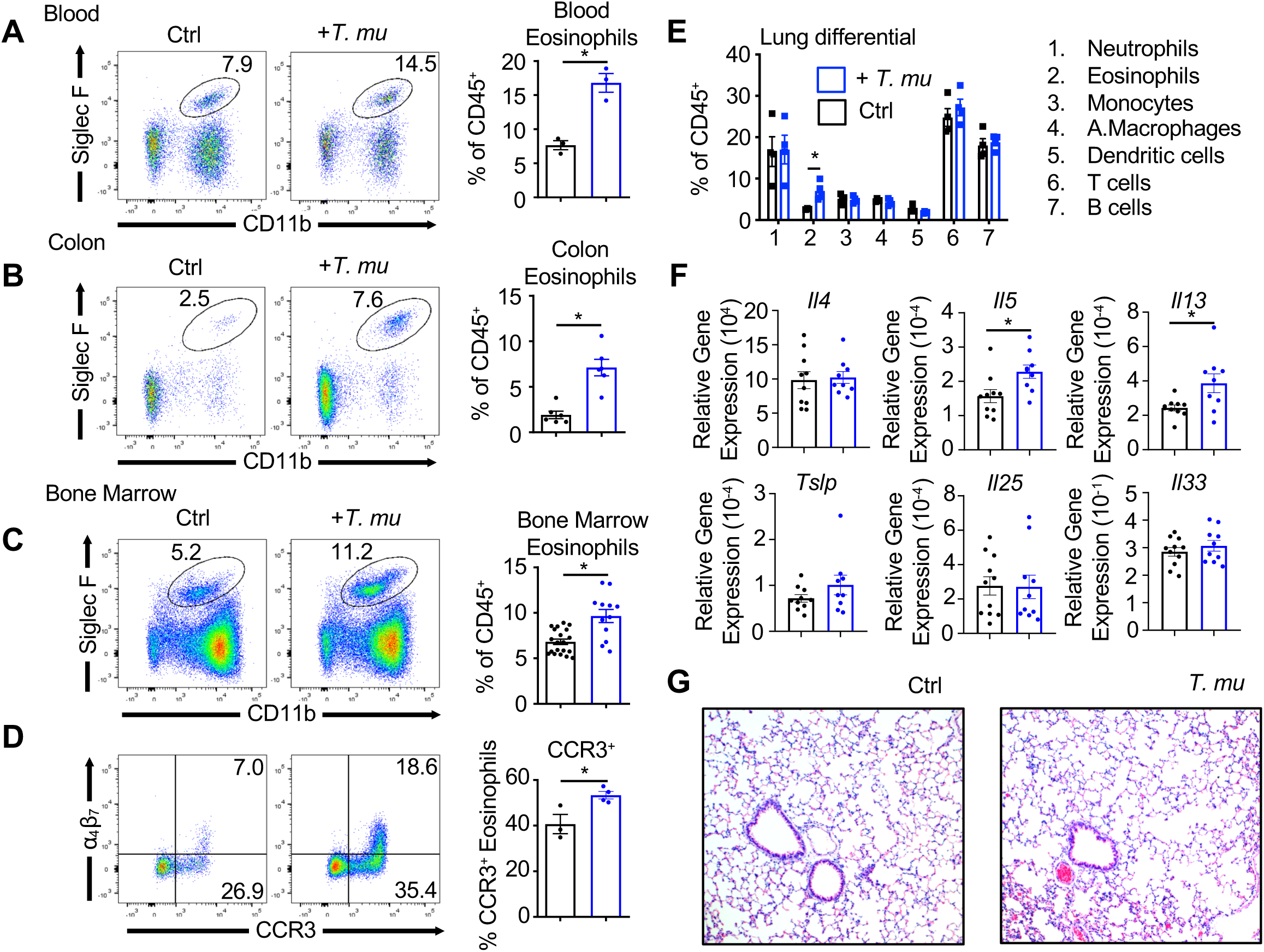
Colonization with *Tritichomonas musculis* elicits systemic eosinophilia. Representative flow cytometry plots and frequencies of eosinophils (CD11b^+^SiglecF^+^) within the CD45+ population of the (A) blood, (B) colon, and (C) bone marrow from naïve and *T.mu*-colonized mice. (D) Frequencies of CCR3 and integrin a4b7 expression on bone marrow eosinophils. (E) Differential analysis of lung immune cells following *T.mu* colonization. (F) Lung mRNA expression of *Il4, Il5, Il13, Tslp, Il25*, and *Il33* relative to *Gapdh*. (G) H&E-stained sections of lungs. Data are shown as mean +/- SEM. Student’s *t* test and one-way analysis of variance (ANOVA) with Bonferroni’s multiple comparison test were performed. Statistical significance is indicated by *P<0.05.

Preferential accumulation of leukocytes at mucosal surfaces requires homing to and interactions with endothelial cells. The integrins α4β7 and the chemokine receptor CCR3 promote mucosal homing of eosinophils (*18, 19*). The BM of *T.mu*^+^ mice contained increased numbers of CCR3^+^α4β7^-^ and CCR3^+^α4β7^+^ eosinophils (Fig.1D). Notably, CCR3 expression has been reported to mediate perivascular accumulation of eosinophils (*20*). CCR3 promotes lung-homing, prompting us to survey all major immune cell types in the lungs of mice colonized with *T.mu*. An almost exclusive increase in eosinophils was observed, while no major changes in other immune cell frequencies were seen (Fig.1E). The accumulation of eosinophils in the lungs of mice carrying the gut protozoan commensal, corresponded to significantly elevated levels of IL-5 and IL-13 mRNA, despite unchanged levels of the type 2 immune activating alarmins thymic stromal lymphopoietin (TSLP), IL-25, and IL-33 (Fig.1F). A histological assessment of the lungs from control and *T.mu*-colonized mice revealed perivascular accumulation of eosinophils while obvious signs of spontaneous inflammation, goblet cell hyperplasia and fibrosis were not observed (Fig.1G and fig.1c and d). These data indicate that colonization by *T.mu* induces a systemic increase in eosinophils with elevated bone marrow eosinophilopoiesis of mucosa-homing eosinophils, demonstrating that a protozoan commensal can regulate tissue-specific and systemic innate type 2 immune cell distribution in healthy mice.

IL-5 and IL-13 are important cytokines for eosinophil survival and are released by tissue-resident ILC2s among other cells (*9*). Lung ILC2s produced higher levels of IL-5 and IL-13 following colonization by *T.mu*, in accordance with higher expression of *Il5* and *Il13* (Fig.1F and Fig.2A). Characterization of lung ILC2s revealed an emerging population of Lin^-^Gata3^+^KLRG1^hi^CD25^-^ iILC2s in the lungs of *T.mu*^+^ mice (Fig.2B). Quantifying iILC2s and nILC2s (the latter defined as: Lin^-^Gata3^+^KLRG1^int/+^CD25^+^) showed a significant increase in iILC2s, but not nILC2 in the lungs of protozoa colonized mice (Fig.2C). In line with previous reports, iILC2s expressed low levels of CD90, CD127 and ST2, but elevated levels of ICOS, IL-17RB, RORγt, and OX40L (fig.2a and b)(*21*). Compared to nILC2s, iILC2s expressed higher levels of *Il5* and *Il13* as well as *Il17a*, coinciding with elevated RORγt levels (Fig.2D, fig.2a and b). Ki67 expression was increased in iILC2s further suggesting that lung-resident iILC2s post colonization with *T.mu* resemble IL-25 elicited gut ILC2s (fig.2a) (*8*). IL-25 injection or helminth infections have previously been shown to expand gut-resident iILC2s and drive interorgan trafficking to the lung in an S1PR1-dependent mechanism (*8*). To confirm that *T.mu*-colonized mice show an elevated bias towards ILC2 activation, mRNA expression for *Il5*, *Il13* and *Il25* were analyzed and found to be increased in contrast to *Il4* or *Il33*, prompting speculations that lung iILC2 may originate from an IL-25 elicited gut-induced pool of iILC2s in *T.mu*^+^ mice (Fig.2E). To determine whether *T.mu*-driven expansion of lung iILC2s is the result of S1PR1-dependent egress, mice colonized with *T.mu* were either left untreated or injected with the S1PR1 antagonist FTY720. Chemical inhibition of the sphingosine receptor completely abrogated the accumulation of iILC2s in the lungs and mesenteric lymph nodes (mLN) but not the intestine after colonization by *T.mu* (Fig.2F). We next sought to address whether antibiotic-mediated eradication of *T.mu* from the intestine could reverse trafficking of iILC2s to the lung and decrease lung eosinophils in *T.mu* colonized mice. Administration of metronidazole, a standard anti-protozoan antibiotic faithfully decreased *T.mu* counts and lowered lung iILC2 and eosinophil numbers (Fig.2G), demonstrating that colonization with the protozoan commensal *T.mu* drives the accumulation of iILC2s in the lungs in S1PR1-depedent fashion.

**Figure 2.**
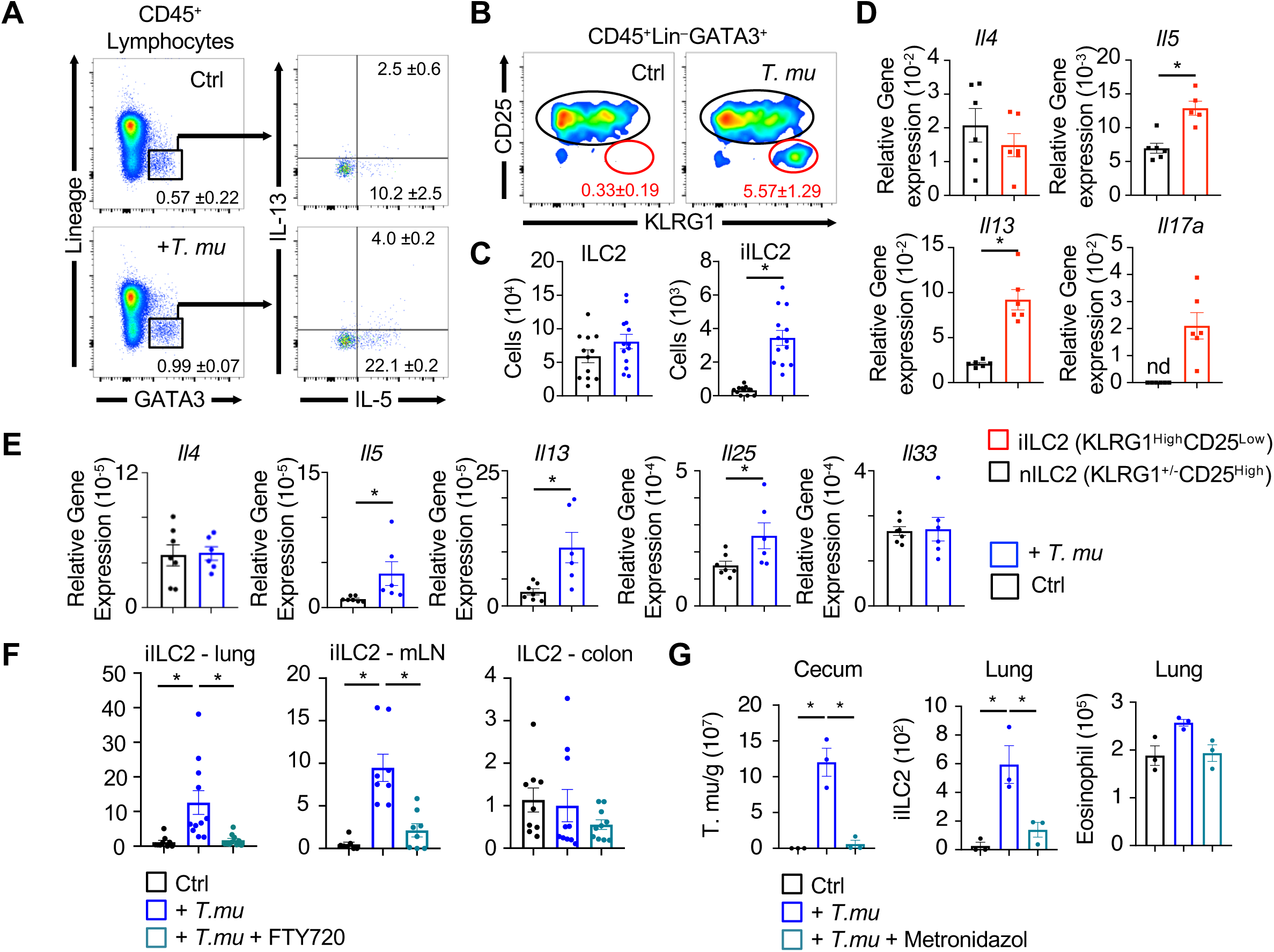
*Tritrichomonas musculis* promotes accumulation of iILC2 in the lung. (A) Representative flow cytometry plots of IL-5 and IL-13 producing live CD45^+^lineage^-^ GATA3^+^ ILC2 in naïve and 21 day *T.mu*-colonized mice. (B) Gating of live CD45^+^lineage^-^ Gata3^+^ ILC2s further gated on CD25^low^ and Klrg1^high^ expression to identify inflammatory ILC2 (iILC2) cells. (C) Enumeration of total lung ILC2 and iILC2. (D) mRNA expression of *Il4, Il5, Il13* and *Il17a* from sort purified lung CD45^+^lineage^-^Gata3^+^CD25^High^Klrg1^+/-^ nILC2 and CD45^+^lineage^-^Gata3^+^CD25^low^Klrg1^High^ iILC2 in *T.mu*-colonized mice. (E) Gut mRNA expression of *Il4, Il5, Il13, Il25,* and *Il33* relative to *Gapdh*. (F) G) Absolute *T.mu* counts in colonized and metronidazole-treated *T.mu*^+^ mice. Adjacent bar graphs show quantification of lung ILC2 and eosinophils in *T.mu*^+^ mice following metronidazole-treatment. Data are shown as mean +/- SEM. Student’s *t* test and one-way analysis of variance (ANOVA) with Bonferroni’s multiple comparison test were performed. Statistical significance is indicated by *P<0.05

We next evaluated whether *T.mu*-mediated expansion of ILC2s drives lung eosinophilia by quantifying eosinophils and iILC2s in ILC-deficient *Rag2^-/-^Il2rg^-/-^* and ILC-sufficient *Rag1^-/-^* mice (Fig.3A). ILC-deficient *Rag2^-/-^Il2rg^-/-^* and ILC2-sufficient *Rag2^-/-^* mice showed no eosinophilia in their lungs (Fig.3A). Interestingly, lung iILC2s in *Rag2^-/-^* mice did not expand following colonization with *T.mu*, in contrast to the observations made in wild type mice (Fig.3A and Fig.2). These observations indicate that Rag1/2-dependent cells participate in the accumulation of both eosinophils and iILC2s in the lung. Analysis of mice lacking either all T cells (*Tcrb^-/-^Tcrd^-/-^*) or B cells (*µMT^-/-^*) revealed that T cells help mediates the accumulation of eosinophils while B cells promote both iILC2 and eosinophil accumulation in the lung following *T.mu* colonization (Fig.3A and B). Adoptive transfer of purified splenic T cells and B cells into *Rag2^-/-^Il2rg^-/-^* mice, was insufficient to reestablish lung eosinophilia in the absence of ILC2s, despite colonization with the protozoan gut commensal (Fig.3C). Transfer of sorted intestinal ILC2s alone, also failed to elevate lung eosinophils even in the presence of *T.mu* (Fig.3C). Only co-transfer of purified intestinal ILC2s together with splenic T cells and B cells into *Rag2^-/-^Il2rg^-/-^* mice restored eosinophilia in the lung following intestinal *T.mu* colonization, revealing a previously unappreciated tripartite interaction between iILC2, T cells and B cells for the induction of lung eosinophils (Fig.3C). Notably, *Rag2^-/-^* mice retained elevated colonic eosinophil counts following colonization with *T.mu*, emphasizing a selective requirement for T cells and B cells in controlling iILC2s and eosinophils in the lung but not the intestinal tract (fig.3a).

**Figure 3.**
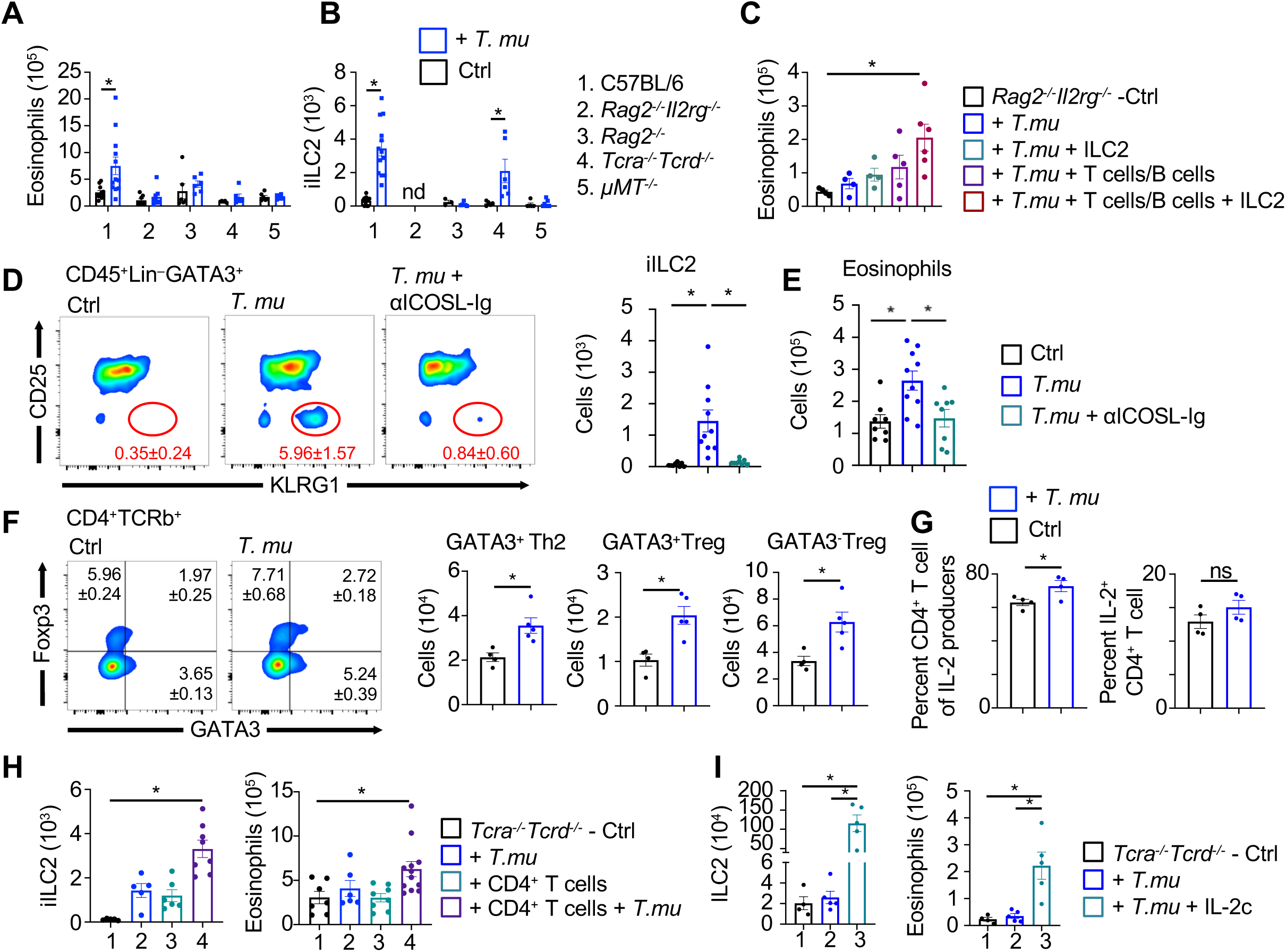
An immune tripartite of B cells, T cells and iILC2s govern lung eosinophilia following colonization by *Tritrichomonas musculis*. Absolute numbers of lung (A) eosinophils and (B) iILC2s from naïve and *T.mu*-colonized C57Bl/6, *Rag2^-/-^Il2rg^-/-^, Rag2^-/-^,Tcra^-/-^Tcrd^-/-^* and *µMT^-/-^* mice. (C) Total lung eosinophil counts from *Rag2^-/-^Il2rg^-/-^*mice colonization with *T.mu* after transfer of T and B cells, gut ILC2, or co-transfer of T cells, B cells and ILC2. (D) Frequencies and absolute counts of lung iILC2 and (E) eosinophils following colonization with *T.mu* and treatment with an anti-ICOSL antibody. (F) Total cell counts and frequencies of lung GATA3^+^Foxp3^-^ Th2, GATA3^+^Foxp3^+^ Treg and GATA3^-^Foxp3^+^Treg cells as well as (G) frequencies of IL-2 producing T cells within the lung of naïve and *T.mu*-colonized C57Bl/6 mice. (H) Total lung iILC2 and eosinophil counts from *Tcra^-/-^Tcrd^-/-^*mice with or without transfer of CD4^+^ T cells or colonization with *T.mu*. (I) Absolute numbers of lung ILC2 and eosinophils in *Tcra^-/-^Tcrd^-/-^*mice colonized with *T.mu* and treated with IL-2c. Data are shown as mean +/- SEM. Student’s *t* test and one-way analysis of variance (ANOVA) with Bonferroni’s multiple comparison test were performed. Statistical significance is indicated by *P<0.05

We next sought to investigate the underlying mechanisms orchestrating this protozoan-driven lymphoid cell network in the airways. Accumulating lung iILC2s expressed higher levels of ICOS compared to nILC2s suggesting the tuning of lung ILC2s through ICOS/ICOSL interactions (fig.2) (*22*). In turn, lung B cells constituted ∼ 90% of all available ICOSL, implicating the engagement of ICOS on iILC2s through B cell-expressed ICOSL as regulator of iILC2 accumulation and the resulting eosinophilia in the lung (fig.3b). Injection of blocking anti-ICOSL antibodies reliably decreased available ICOSL on CD45^+^ lung cells and reverted the *T.mu*-driven accumulation of iILC2s and eosinophils in the lung, indicating that B cell-mediated engagement of ICOS on iILC2s serves as critical regulator of lung iILC2s and eosinophils (fig.3c, Fig.3D and E). *Tcrb^-/-^Tcrd^-/-^* mice colonized with *T.mu*, failed to show an elevation in lung eosinophils, indicating the involvement of T cells in the *T.mu*-driven immune adaptation of the lung (Fig.3A and B). Lung-resident iILC2s express OX40L, an activator of OX40-expressing CD4^+^ T cells. Engagement of OX40-OX40L increases the production of IL-2 and Th2-associated IL-5 possibly adding to accumulation of eosinophils in the lung (*23*). Interestingly, lung eosinophilia has been reported to be a side effect of IL-2:anti-IL-2-Ig immune complex treatment (IL-2c), an IL-2 receptor agonist therapy used to expand regulatory T (T_reg_) cells (*24*). *T.mu*-colonized mice showed a significant increase in Foxp3^-^ Gata3^+^ Th2 cells, as well as Gata3^+^ and Gata3^-^ Foxp3^+^ T_reg_ cells in their lung, suggesting a higher availability of IL-2 (Fig.3F). Quantification of IL-2 production in control *or T.*mu^+^ mice revealed CD4^+^ T cells as major source for steady state IL-2 and elevated level thereof in the presence of the protist (Fig.3G and fig.3d). T cell-derived IL-2 supports anti-helminth immunity through the activation of ILC2s (*25*). To determine whether IL-2 producing CD4^+^ T cells constitute a regulator of pulmonary iILC2s and thus eosinophils, purified splenic CD4^+^ T cells were injected into *T.mu* colonized *Tcrb^-/-^Tcrd^-/-^* mice. Engraftment of CD4^+^ T cells in *Tcrb^-/-^Tcrd^-/-^* mice increased lung iILC2s and eosinophil numbers confirming the requirement of CD4^+^ T cells in executing *T.mu*’s remote regulation of lung immune cells (Fig.3H). To investigate whether IL-2 production by T cells could restore lung eosinophilia, *Tcrb^-/-^Tcrd^-/-^* mice were injected with IL-2c immunocomplexes. T cell-deficient mice, colonized with *T.mu*, accumulated ILC2s and eosinophils only after injection of IL-2c, establishing that T cell-derived IL-2 regulates lung innate immune cells following colonization with *T.mu* (Fig.3I). Collectively these findings demonstrate that CD4^+^ T cells mediate the accumulation of lung eosinophils after colonization with the gut commensal protist *T.mu* in an IL-2 dependent manner.

Increased ILC2 activation and elevated numbers of eosinophils are a feature of allergic airway inflammation at least in a subgroup of patients. Our observation of elevated iILC2 and eosinophil numbers in the lung of *T.mu*-colonized mice prompted us to determine whether the protozoan-driven remote reprogramming of the lung immune landscape could modulate the outcome of allergic airway inflammation. Inhalation of house dust mite (HDM) extract in mice sensitizes towards the development of allergic asthma, which manifests as increased infiltrating hematopoietic cells, including eosinophils, elevated serum immunoglobulin (Ig) E levels, goblet cell hyperplasia and collagen deposition, recapitulating the major pathologies of human asthma (Fig.4A and B, fig.4A-C). Conventional C57BL/6 mice reproducibly developed HDM-driven airway inflammation with anticipated histopathological scores (Fig.4A and B, fig.4A-C). Animals colonized with *T.mu* showed significantly higher pathological scores, particularly alveolar and interstitial inflammation reached scores beyond those commonly observed for this model (Fig.4A and B, fig.4A-C). Goblet cell hyperplasia and collagen deposition did not exceed those recorded for conventional mice (fig.4A-C). In line with these observations, bronchioalveolar lavages isolated from HDM-treated, *T.mu*-colonized mice contained significantly higher numbers of CD45^+^ cells, dominated by eosinophils (Fig.4C). The disease promoting capability of eosinophils supports the histopathologic observations of exacerbated allergic airway inflammation in *T.mu*^+^ mice, demonstrating that protozoan-driven remote regulation of the lung immune system increases the severity of allergic airway inflammation through a gut-driven, lung-resident immune network between T cells, B cells and iILC2s.

**Figure 4.**
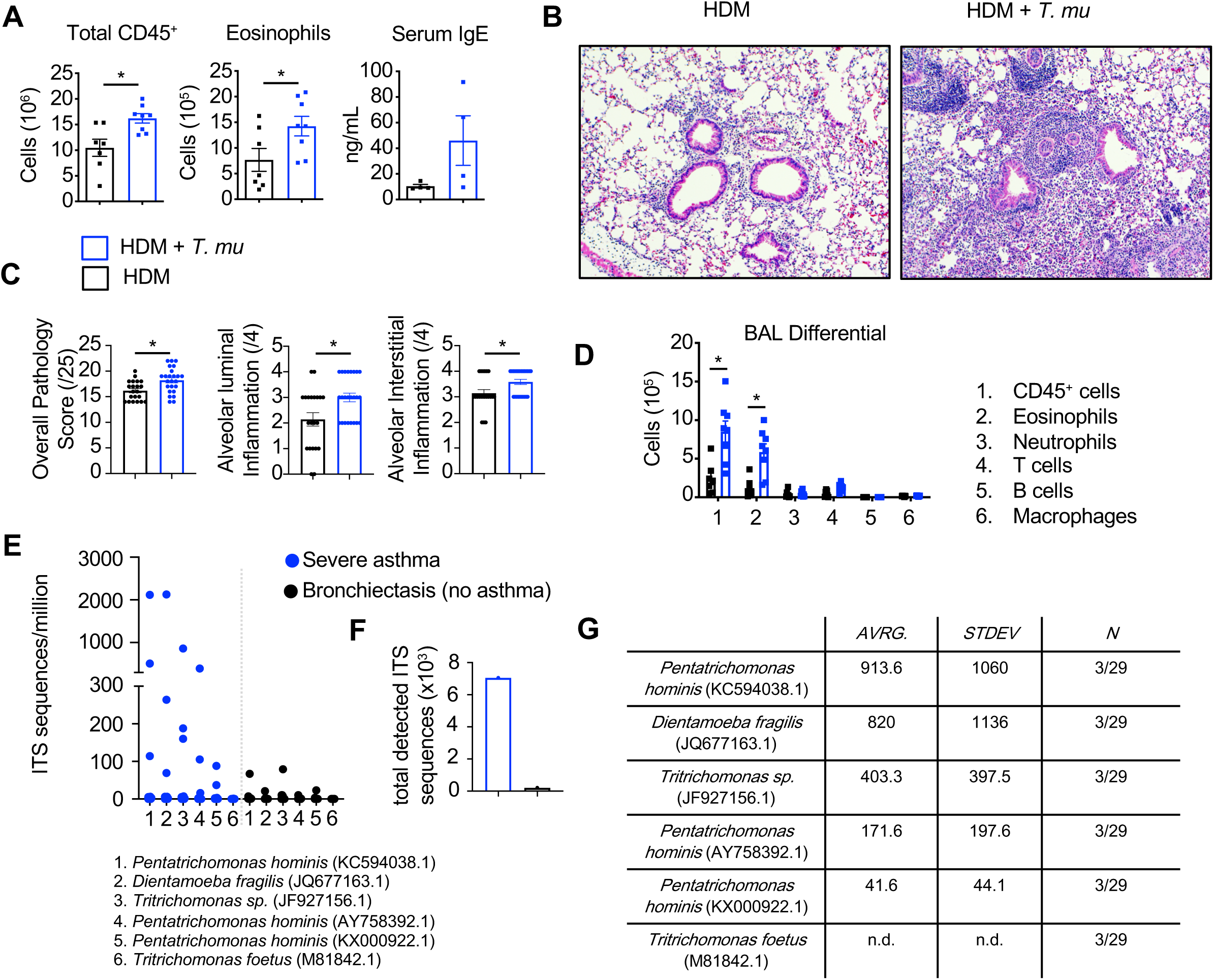
Protozoan commensals exacerbate allergic airway inflammation and are detectable in asthma patients. (A) Quantification of lung CD45^+^ cells and eosinophils as well as serum IgE following house dust mite (HDM)-induced allergic airway inflammation. (B) H&E-stained sections of lungs. (C) Overall pathology and alveolar luminal and interstitial inflammation scores from lung histological samples. (D) Bronchioalveolar lavage (BAL) fluid immune cell differential and total counts. (E) Survey for protozoan ITS sequences in metagenomic DNA samples from severe asthmatic (SA) and bronchiectasis (BE) patients. Dot plot shows numbers of protozoa-specific ITS sequences. (F) Quantification of all selected protozoan ITS in SA and BE patients. (G) ITS sequence counts and variability across analyzed patients.

*Dientamoeba fragilis* and *Pentatrichomonas hominis* are two human-associated protozoan species, closely related to *T.mu*, based on their similarity in the internal transcribed spacer (ITS) sequences (*12*). A systemic survey of protozoan ITS sequences in patients diagnosed with allergic airway inflammation has yet to be produced and is currently not available. Interestingly, a previous case report study implicated the presence of protozoa in the sputum and airways of patients diagnosed with asthma (*26*). Even though these observations remain controversial, they suggest a possible involvement of protozoa in human allergic airway inflammation. To determine whether traces of protozoan DNA are detectable in the sputum of patients with allergic airway inflammation, we made use of a publicly available metagenomic dataset (*27*). Sequences were previously generated through shot-gun sequencing of DNA isolated from pulmonary samples derived from severe asthma patients (SA, n=29) and non-asthmatic patients diagnosed with bronchiectasis (BE, n=14). Bronchiectasis is a condition accompanied by a thickening of the airway wall, primarily mediated by chronic inflammation. We performed an informed analysis of the publicly available metagenomic dataset from SA and BE patients for ITS sequences of human-associated *Tritrichomonas* spp., *Dientamoeba fragilis* and *Pentatrichomonas hominis*. ITS sequences specific for the cattle and cat-associated protist *Tritrichomonas foetus* were used as a negative control. Indeed, neither BE, nor SA patients showed ITS sequences specific for *Tritrichomonas foetus* confirming the specificity of the analysis (Fig.4E). SA patients identified with ITS sequences showed an overall greater abundance of ITS sequences specific for the human-related protozoan species when compared to the non-asthmatic BE control group (Fig.4F). These findings demonstrate that protozoan traits are detectable in human isolates from patients diagnosed with severe asthma but remain moderate, as the number of SA patients showing protozoa-specific ITS sequences constituted ∼10% (Fig.4G). However, these 10% nevertheless presented with significantly elevated copies of ITS sequences (Fig.4F and G). The high variability in the frequency of ITS sequences across SA patients, indicate a possible underlying heterogeneity within protozoa^+^ SA patients or inconsistency in sample collection. These results indicate that a subgroup of severe asthma patients show signs of protozoan colonization and may possibly constitute a new subset of asthma patients (Fig.4E-G). Future validation in larger and better controlled cohorts is needed to confirm these findings and establish a possible new sub-classification of asthma patients based in the statues of their commensal protozoan colonization.

Asthma is known to be influenced by the gut microbiota with states of early dysbiosis increasing the risk for the development of disease in humans and mice. The focus of these studies targeted bacterial communities within the microbiota, which limit the depth of these studies and their ability to demonstrate causal connections between specific microbes and the resulting disease (*4*). Our data demonstrate that the gut commensal protozoa *T.mu* drives a remote regulated lung-specific tripartite immune network, resulting in the accumulation of eosinophils, which in turn exacerbate the severity of allergic airway inflammation. Protozoan-induced IL-25 production elicits iILC2s in the intestine and facilitate an S1PR1-dependent interorgan migration of these cells into the lung. Accumulation of iILC2s in the lung critically relied on support from lung-resident CD4^+^ T cells and B cells to facilitate eosinophilia in this organ. T cell-derived IL-2 and B cell-expressed ICOSL were required signals provided by lung-resident adaptive immune cells to sustain the accumulation of migratory innate type 2 cells. Adoptive co-transfer of gut-derived iILC2s, splenic T cells and B cells were a requirement to reinstate lung eosinophilia following colonization with *T.mu*. In turn, blockade of ICOSL or treatment with IL-2c confirmed the underlying molecular interactions of this new tripartite immune network. The resulting increase in eosinophils following the ignition of lung iILC2s - T cell - B cell interactions exacerbated allergic airway inflammation in protozoa-colonized mice. Collectively, these results demonstrate that the accumulation of eosinophils in the lung can be regulated across organs and requires a specific local network of cells to manifest a local asthma-promoting niche for eosinophils.

In patients, asthma is a heterogeneous disease promoted by several underlying mechanisms. Therapeutic options follow a rigid step-wise approach which demands in severe cases, trial and error approaches in case of a lack of symptom control. Thus, multiple treatment attempts can be necessary prior to achieving an optimal therapeutic management (*28*). Intriguing investigations of serological biomarkers allow the categorization of patients into subclasses and confirmed the effectiveness of such approaches to tailored intervention (*29*). We report that protozoan colonization promotes lung eosinophilia in an ILC2-dependent manner with minimal elevations of IL-13 driven remodeling of the tissue, suggesting a distinct pathology. Observing protozoan DNA sequences in a small, yet sizeable group of human pulmonary isolates suggests that a microbiota-focused classification of patients, based on protozoan traits, may result in a new sub-classification of patients’ endotypes. Our analysis of the underlying immunological pathways responsible for the protozoa-driven immune adaptation in the lung provides multiple cellular and molecular targets that may proof useful in a putative subgroup of asthma patients.

The newly uncovered role for gut protozoan commensals in the threshold for disease-associated immune cells in the lung has broad implications in understanding how the intestinal microbiota, beyond their bacterial communities, initiates gut-tissue axes to remotely shape the immune landscape of distinct organs. These mechanisms involve previously unknown modes of innate immune regulation by tissue-resident adaptive immune cells and represent a significant advancement in our understanding of protozoa as gut commensal regulators of host immunity. Their remote regulation of the host’s immune system may provide insight for the development of new diagnostic approaches as well as new therapeutics for the treatment of allergic airway inflammation in a new sub-class of asthma patients.

## Acknowledgments

We would like to thank all members of *#theonlylabever* for their support and discussion. We acknowledge the excellent support by the University of Toronto, Temerty Faculty of Medicine Flow Cytometry Core facility. We wish to thank Dr. Thomas Eiwegger, Dr. Juan-Carlos Zúñiga-Pflücker and Dr. Jennifer Gommerman for critical reading of the manuscript.

## Funding

CIHR Banting Postdoctoral Fellowship Program (KB)

Vanier Canada Graduate Scholarship - NSERC (PC)

Canada Graduate Scholarships - Doctoral program (LN)

Canadian Institutes for Health Research - PJT-388337 (AM)

Canadian Institutes for Health Research – 6210100847 (AM)

JP Bickell Foundation – (AM)

Natural Sciences and Engineering Research Council - RGPIN-2019-04521 (AM)

Tier 2 Canada Research Chair Program (AM)

Science Foundation Ireland Centre Grant – 12/RC/2273_P2 (LOM)

## Author contributions

Conceptualization: KB, AM

Methodology: KB, CS, BF, LOM, AM

Investigation: KB, PC, LN, EC, CS, BF, LOM, AM

Visualization: KB, BF, LOM, AM

Funding acquisition: LOM, AM

Project administration: AM

Supervision: AM

Writing – original draft: KB, AM

Writing – review & editing: KB, AM

## Competing interests

“Authors declare that they have no competing interests.”

## Data and materials availability

“Data and material will be available in the main text or the supplementary materials.”

## Supplementary material

### Material and Methods

#### Mice

C56Bl/6, B6.129P2-Tcrb^tm1Mom^ Tcrd^tm1Mom/J^ (*Tcrb^-/-^Tcrd^-/-^)*, and B6.129S2-Ighm^tm1Cgn/J^ (*µMT^-/-^)* mice were purchased from Jackson Laboratory while B6.129S6-Rag2^tm1Fwa^ *(Rag2^-/-^)* and C57BL/6NTac.Cg-*Rag2^tm1Fwa^ Il2rg^tm1Wjl^ (Rag2^-/-^Il2rg^-/-^)* mice were purchased from Taconic. Mice were subsequently bred in-house under specific pathogen-free conditions at the University of Toronto, Division of Comparative Medicine. All animal procedures were conducted using age- and sex-matched littermates and were approved by the University of Toronto Animal Care Committees and performed in accordance with the committees’ ethical standards.

#### Colonization with Tritrichomonas musculis

The cecal content of *T. mu* colonized mice was harvested into 20ml of sterile PBS and filtered several times through a 70μm cell strainer. The suspension was pelleted, and the supernatant was discarded. *T.mu* were purified from the pellet by centrifugation with a 40/80% Percoll (GE Healthcare Life Sciences) gradient and were collected from the interface. *T.mu* was further purified via sorting on a FACS ARIA IIIu. Two million purified *T. mu* were gavaged per recipient mouse immediately following the sort.

#### House dust mite model of allergic airway inflammation

Mice were anesthetized under aerosolized isoflurane and intranasally instilled with 100 μg of house dust mite (HDM) antigen (Greer, Lenoir, NC) for 3 consecutive days (days 0, 1, and 2). On days 13 to 17 post sensitization, mice were intranasally challenged with 25 μg of HDM antigen daily. In some cases, mice were colonized with *T.mu* 10 days before the first HDM sensitization. On day 18, mice were injected intraperitoneally (i.p.) with 2,2,2-tribromoethanol (Avertin; Sigma), tracheas were cannulated, and bronchoalveolar lavages (BALs) were performed using triplicates of 1 ml of sterile 10% fetal bovine serum in PBS. BAL fluid was then red cell lysed using ammonium chloride buffer.

#### Isolation of leukocytes from tissue

Colon leukocytes were isolated as previously described (*1*). Briefly, colon tissue was cut open longitudinally and then briefly washed with ice-cold phosphate-buffered saline (PBS). Tissue was incubated in Hank’s balanced salt solution (HBSS) without calcium/magnesium (Gibco) containing 10 mM EDTA for 15 min at 37 °C and extensively vortexed to remove epithelial cells. Tissue was then digested with 0.5mg/mL Collagenase (Sigma) and 0.5mg/mL DNase I (Sigma) in HBSS on a shaker at 100 rpm, 37 °C, for 20 min, extensively vortexed and filtered through a 70μm cell strainer. Cells were purified by centrifugation with a 40/80% Percoll (GE Healthcare Life Sciences) gradient and leukocytes were collected from the interface.

Lung tissues were harvested after perfusion and were chopped into fine pieces then digested for 20 min at 37 °C with 0.5mg/mL Collagenase and 0.5mg/mL DNase I in HBSS. Tissue pieces were sheared through a 18g needled and passed through a 70mm cell strainer. Leukocytes were purified by centrifugation with 40/80% Percoll gradient.

Bone marrow was isolated from a single femur clean of any muscle or other tissues. The ends of bone were cut off with scissors and a 27-gauge needle was used to flush out the marrow with PBS. The marrow was then passed through a 70 μm cell strainer to obtain single-cell suspensions and samples were subsequently red cell-lysed using ammonium chloride buffer.

#### Flow cytometry and cell sorting

Absolute numbers of cells were determined via hemocytometer. Intracellular cytokine staining was performed by stimulating cells with phorbol 12-myristate 13-acetate, ionomycin, and protein transport inhibitor cocktail containing Brefeldin A and Monensin (eBioscience) for 4 h and fixing/permeabilizing cells using the eBioscience intracellular cytokine buffer kit. All antibody dilutions and cell staining were done with PBS containing 2% fetal calf serum, and 5 mM EDTA. Fixable Viability Dye eFluor 506 (eBioscience) was used to exclude dead cells from analyses. Prior to staining, samples were Fc-blocked with buffer containing anti-CD16/32. Samples were either analyzed on an LSR Fortessa X-20 (BD) or sorted with a FACS ARIA IIIu (BD).

#### Antibodies and reagents

Cells were stained with fluorescent conjugated: CD3 (145-2C11; Biolegend), B220 (RA3-6B2; Biolegend), CD25 (PC61.5; eBioscience), NK1.1 (PK136; Biolegend), CD127 (A7R34; Biolegend), a4b7 (DATK32; Biolegend), IL-13 (ebio13a; eBioscience), IL-5 (TRFK5; Biolegend), CD90 (53-2.1; Biolegend), ICOS (C398.4A; Biolegend), CD117 (ACK2; Biolegend), ST2 (RMST2-2; eBioscience), KLRG1 (2F1; eBioscience), Sca1 (D7; Biolegend), CD4 (GK1.5; Biolegend), CD8 (53-6.7; Biolegend), CD45.2 (104; Biolegend), IL-17RB (9B10; Biolegend), Siglec F (1RNM44N; eBioscience), CD11b (M1/70; eBioscience), CD125 (T21; BD), CD11c (N418; Biolegend), Gr1(RB6-8C5; eBioscience), MHCII(M5/114.15.2; Biolegend), Ly6C (HK1.4; eBioscience), RORgt (B2D; eBioscience), OX40L (RM134L; eBioscience), Foxp3 (MF-14; eBioscience), ICOSL (HK5.3; Biolegend), IL-2 (JES6-5H4;Biolegend)

#### Adoptive transfer of lymphocytes

ILC2 were sorted from the lung and small intestine of donor C57Bl/6 mice and injected intravenously into *Rag2^-/-^Il2rg^-/-^* recipient mice. One week later, T cells and B cells were sorted from the spleen of C57Bl/6 mice and injected intravenously into the same *Rag2^-/-^ Il2rg^-/-^* recipient mice. Two weeks later, recipient mice were colonized with 2×10^6^ T.mu by oral gavage. Lung tissue was analyzed 3 weeks post *T.mu* inoculation.

#### Adoptive transfer of CD4 T cells

2×10^5^ CD4^+^CD45RB^high^CD25^-^ were sorted from the spleen of C57Bl/6 mice and injected i.p. into *Tcrb^-/-^Tcrd^-/-^* recipient mice. Two weeks later, recipient mice were colonized with 2×10^6^ *T.mu* by oral gavage. Lung tissue was analyzed 3 weeks post *T.mu* inoculation.

#### In vivo blockade of ICOSL

Mice were infected with *T. mu* as described above. On day 0 post infection, mice were injected i.p. with 500 μg of either control IgG1 or anti-ICOSL (CD275) (BioXcell) diluted in sterile PBS. Mice were repeatedly injected thereafter 3 times per week prior to sacrifice on day 21.

##### FTY720 Treatment

*T.mu* colonized mice were injected i.p. with 1 mg/kg of FTY720 (Selleck Chemicals) constituted in sterile PBS on the day of infection and thereafter injected 3 times per week prior to sacrifice on day 21 post infection.

#### IL-2 Complex Administration

IL-2 complexes were prepared by incubating IL-2 (Peprotech) and anti-IL-2 mAb (JES6; BioXcell) at a 1:5 w/w ratio for 20 min at 37°C. Complexes were then diluted in PBS and administered i.p. (1 μg IL-2 and 5 μg anti-IL-2 per dose) three times per week to day 21 post *T.mu* colonization.

#### RNA isolation and quantitative real-time PCR

Cells were sorted directly into lysis buffer and RNA was extracted using the using RNeasy Mini kit (Qiagen) according to the manufacturer’s instructions. cDNA was generated using High-Capacity cDNA reverse transcription kits (ThermoFisher Scientific). Quantitative PCR was performed using PowerUp SYBR green master mix (ThermoFisher Scientific) and SYBR green-optimized primer sets run on an CFX384 real-time PCR machine (Biorad). Cycle threshold (C_T_) values were normalized relative to *Gapdh* gene expression. The primers used were synthesized *de novo*:

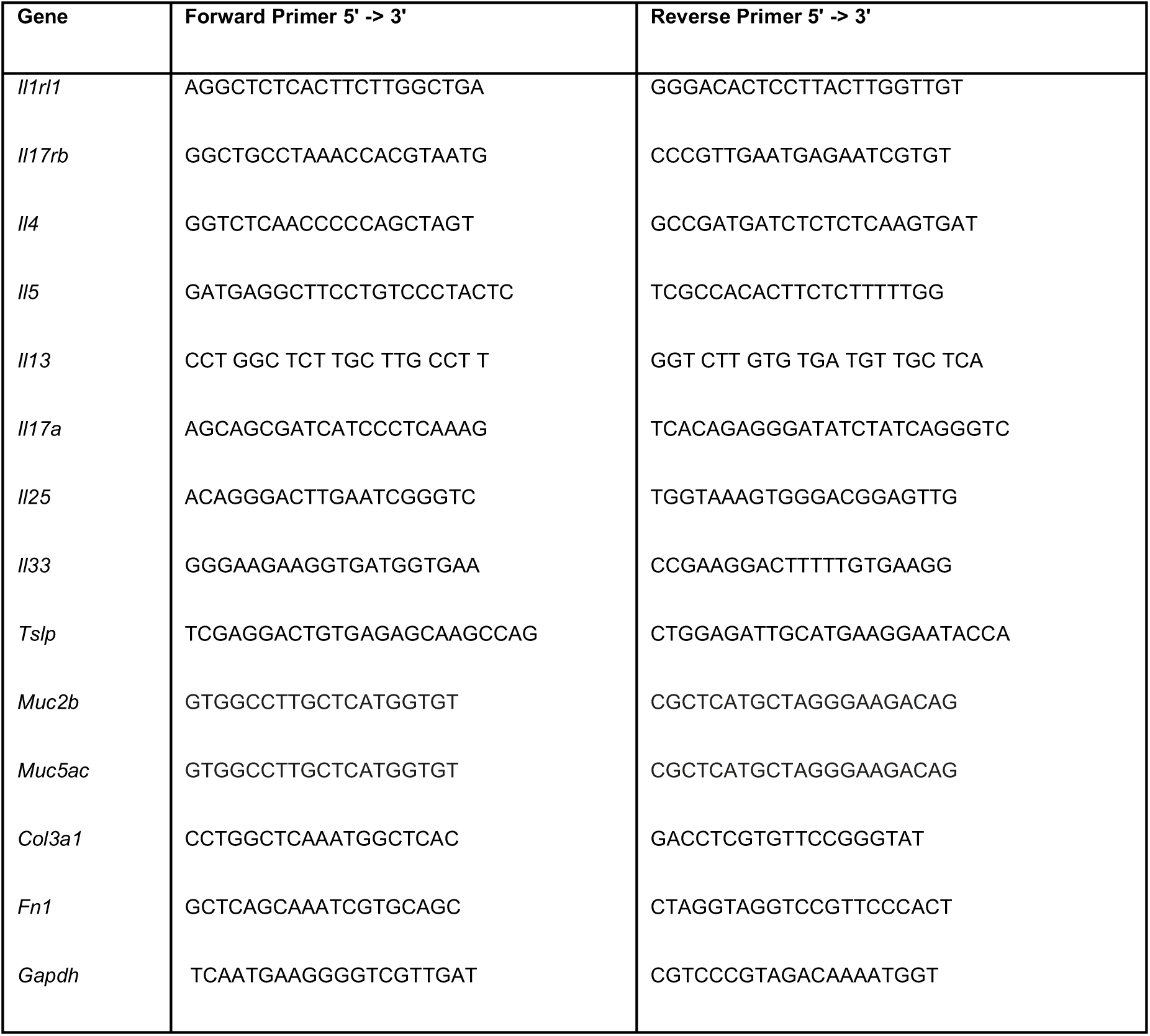

#### ITS sequence analysis

ITS sequences were searched for in a previously published metagenomic dataset associated with “Metagenomics Reveals a Core Macrolide Resistome Related to Microbiota in Chronic Respiratory Disease” (*2*). The SRA files that contained the data were downloaded and transformed into fastq files through the use of the SRA-toolkit as well as the related peripheral “parallel-fastq-dump”. The tool BBDuk was then used to ensure the quality of the reads and the tool Bowtie2 was used to remove host sequences with the hg19 human genome assembly database as a reference. The resulting fastq files were then converted into fasta files through the use of the command line and following this blastn was ran on the files using the ITS sequences as the reference sequences.

#### Histology

Lung tissues were fixed overnight in 10% buffered formalin and paraffin embedded. A total of 5-μm-thick tissue sections were stained with hematoxylin and eosin (H&E), periodic-acid Schiff (PAS), or Masson’s Trichrome for histological analysis.

#### Histology Scoring

Lung sections were scored by a blinded pathologist according to the scoring criteria below.

**(1)** perivascular edema (0, absent; 1, mild to moderate, involving fewer than 25%of the perivascular spaces; 2, moderate to severe, involving more than 25% but less than 75% of perivascular spaces; or 3, severe, involving more than 75% of perivascular spaces);
**(2)** perivascular/peribronchial acute inflammation (0, absent; 1, mild acute inflammation in the perivascular edematous space, with fewer than 5 neutrophils per high-power field; 2, moderate acute inflammation in the perivascular spaces, extending to involve the peribronchial spaces, with more than 5 neutrophils per hpf in these regions; or 3, severe, acute inflammation in the perivascular and peribronchial spaces with numerous neutrophils encircling most [50%] of bronchioles);
**(3)** goblet-cell metaplasia of bronchioles (0, absent; 1, few goblet cells present in one or two bronchiolar profiles; or 2, large numbers of goblet cells present);
**(4)** eosinophilic macrophages in alveolar spaces (0, absent; 1,present in fewer than 25% of alveolar spaces; or 2, present in 25% of alveolar spaces).

A total inflammatory score (range 0 to 10), taken as the sum of the individual scores, was determined.

---

**Supplemental Figure 1.**
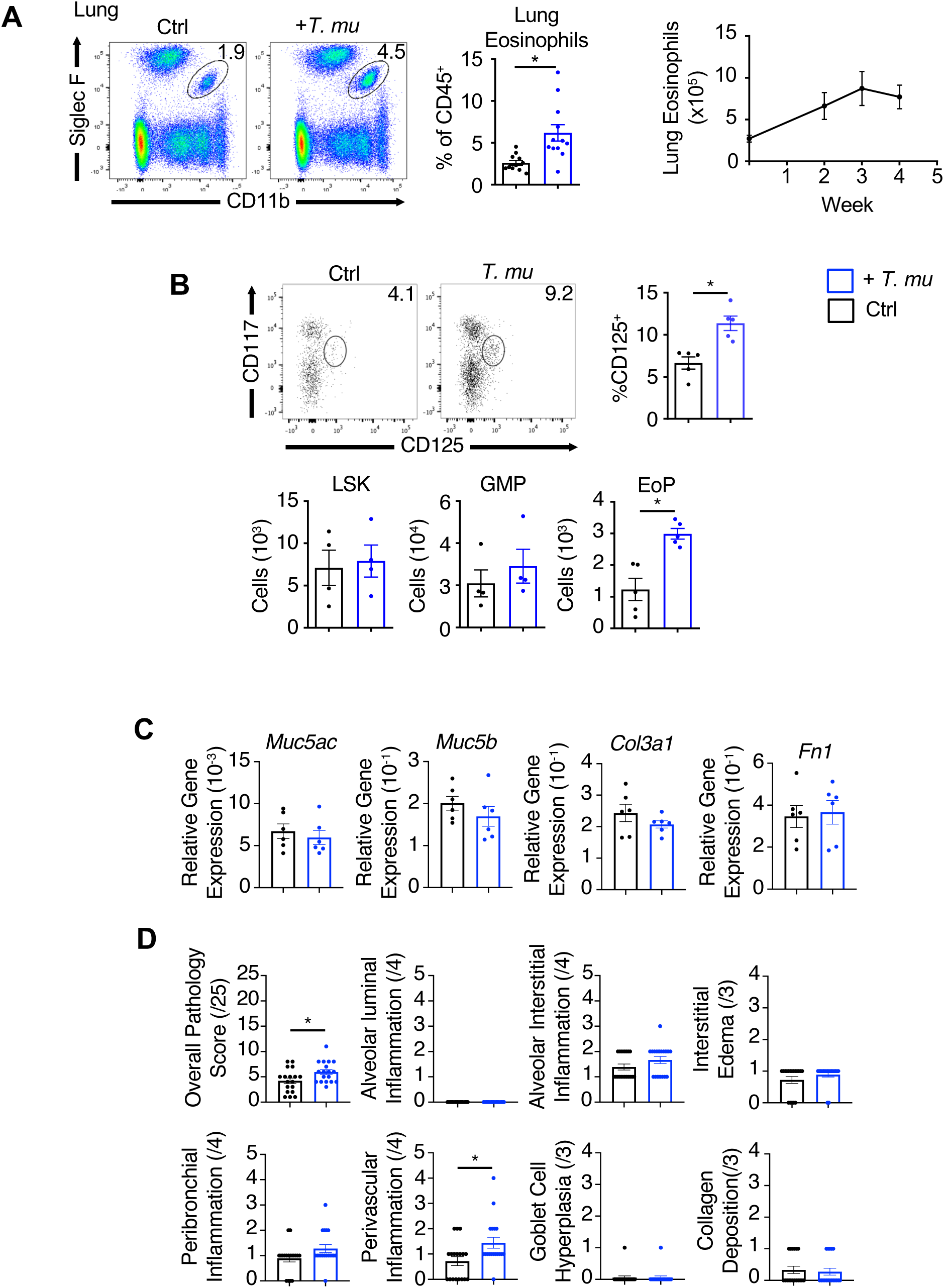
Elevated bone marrow eosinophilopoiesis following colonization by *Tritrichomonas musculis*. (A) Representative flow cytometry plots and frequencies of lung eosinophils 21 days following *T.mu* colonization as well as enumeration of absolute counts of lung eosinophils over the course of T.mu colonization. (B) Frequencies and/or total LSK(lineage^-^ Sca1^+^ckit(CD117)^+^) cell, Granulocyte/monocyte progenitor (GMP; lineage^-^ ckit^+^CD16/32^+^CD34^+^) and eosinophil progenitor (EoP; lineage^-^Sca1^-^ CD34^+^ckit^intermediate^CD125^+^) counts from bone marrow. (C) Lung mRNA expression of *Muc5ac, Muc5b, Col3a1* and *Fn1* relative to *Gapdh*. (D) Pathological scores from analysis of histological lung sections stained with H&E, PAS, and Masson’s Trichrome. Data are shown as mean +/- SEM. Student’s *t* test was performed; statistical significance is indicated by *P<0.05.

**Supplemental Figure 2.**
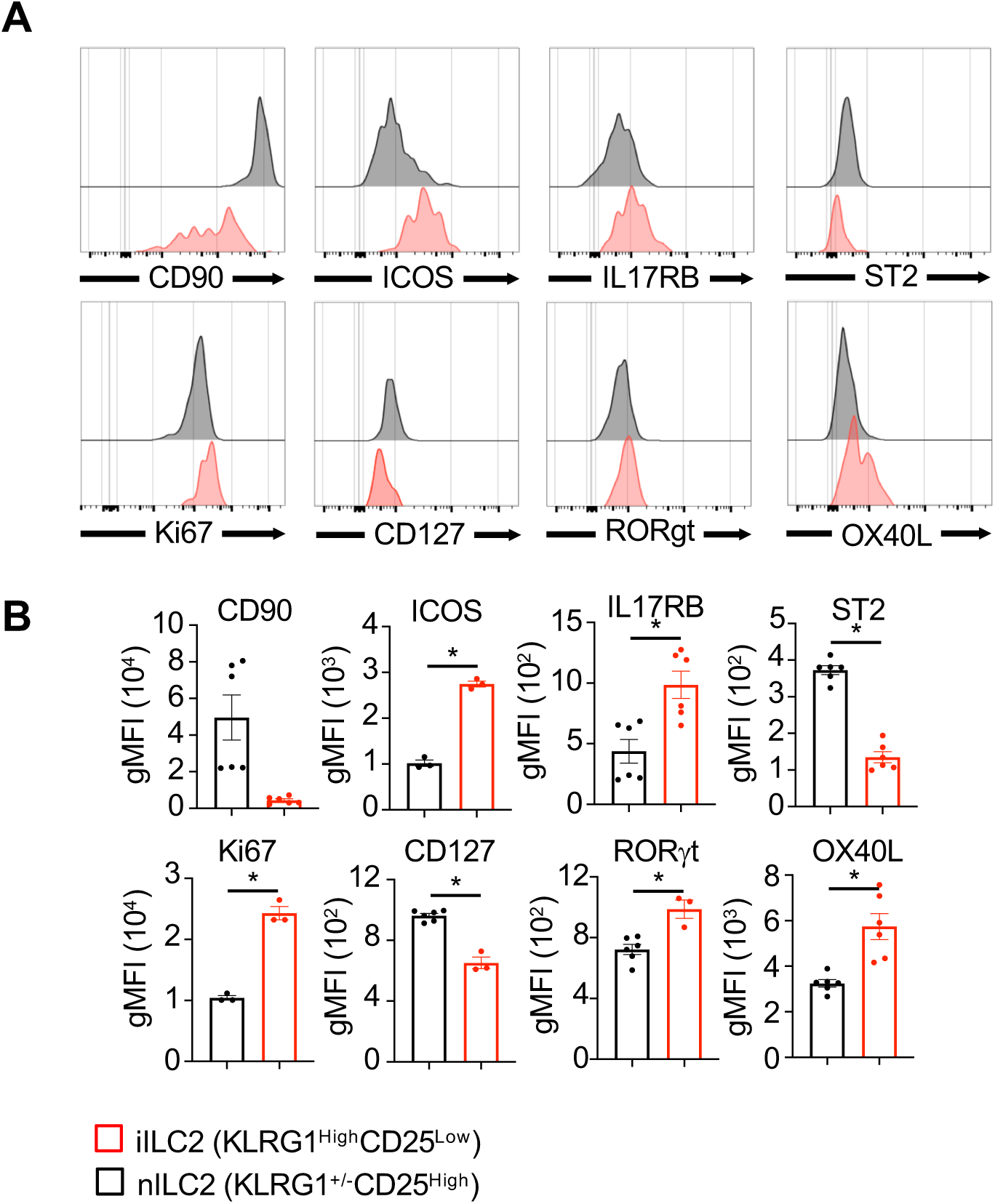
Inflammatory ILC2s are distinct lung ILC2s in *Tritrichomonas musculis*-colonized mice. (A) Representative flow cytometry histograms and (B) quantified geometric mean fluorescent intensity (gMFI) for CD90, ICOS, IL17RB, ST2, Ki67, CD127, RORγt, and OX40L for lung nILC2 and iILC2 from T.mu-colonized mice. Data are shown as mean +/- SEM. Student’s *t* test was performed; statistical significance is indicated by *P<0.05.

**Supplemental Figure 3.**
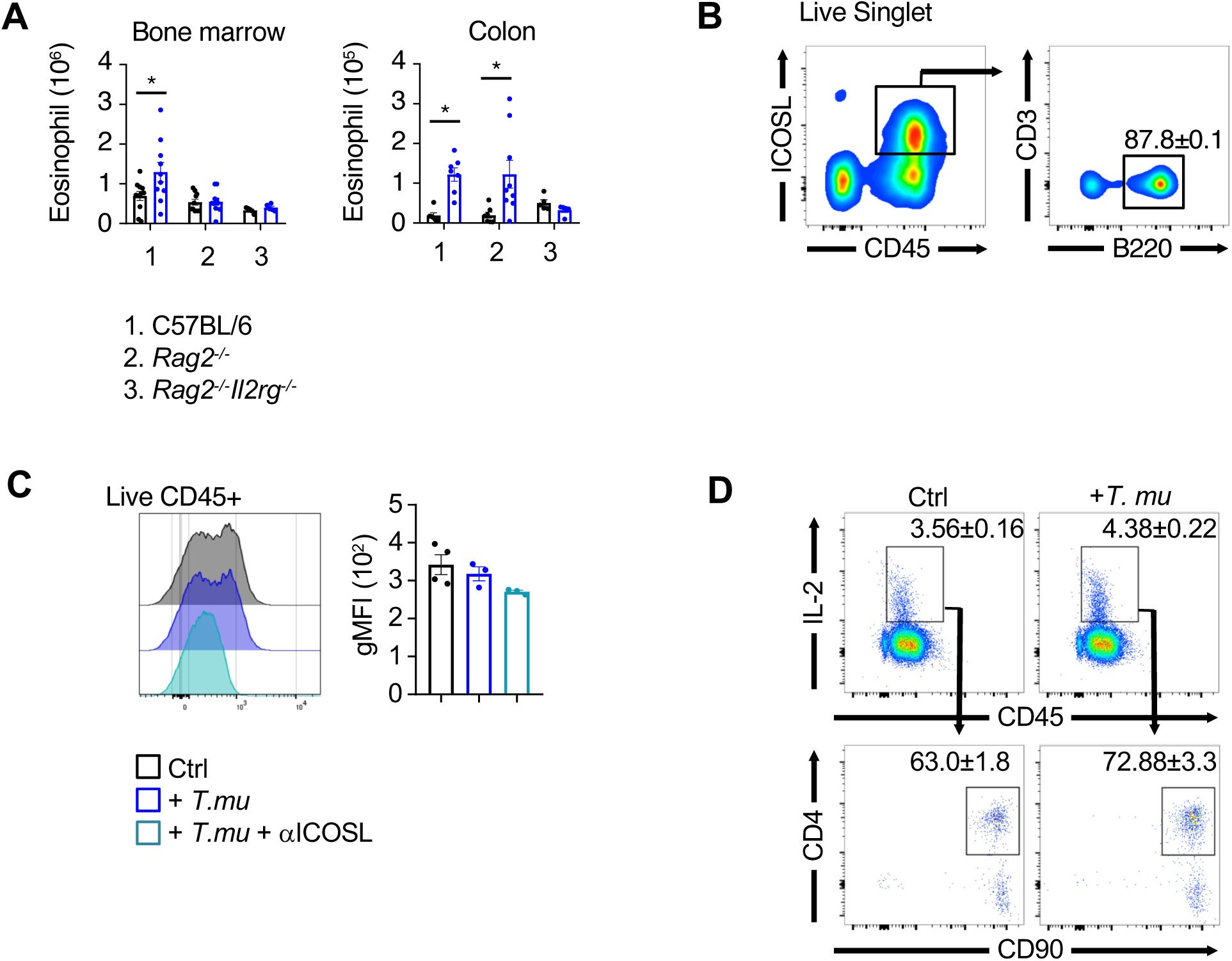
Identification of ICOSL- and IL-2 expressing cells in the lung of *Tritrichomonas musculis*-colonized mice. (A) Bone marrow and colonic eosinophil numbers in untreated of *T.mu*-colonized mice. (B) Gating of ICOSL expressing lung cells. (C) Flow cytometry analysis of ICOSL expression and blockade on immune cells within the lung following *T.mu* colonization +/- treatment with an anti-ICOSL antibody. (D) Gating of IL-2 expressing lung cells in naïve and *T.mu* colonized mice. Data are shown as mean +/- SEM. One-way analysis of variance (ANOVA) with Bonferroni’s multiple comparison test was performed. Statistical significance is indicated by *P<0.05

**Supplemental Figure 4.**
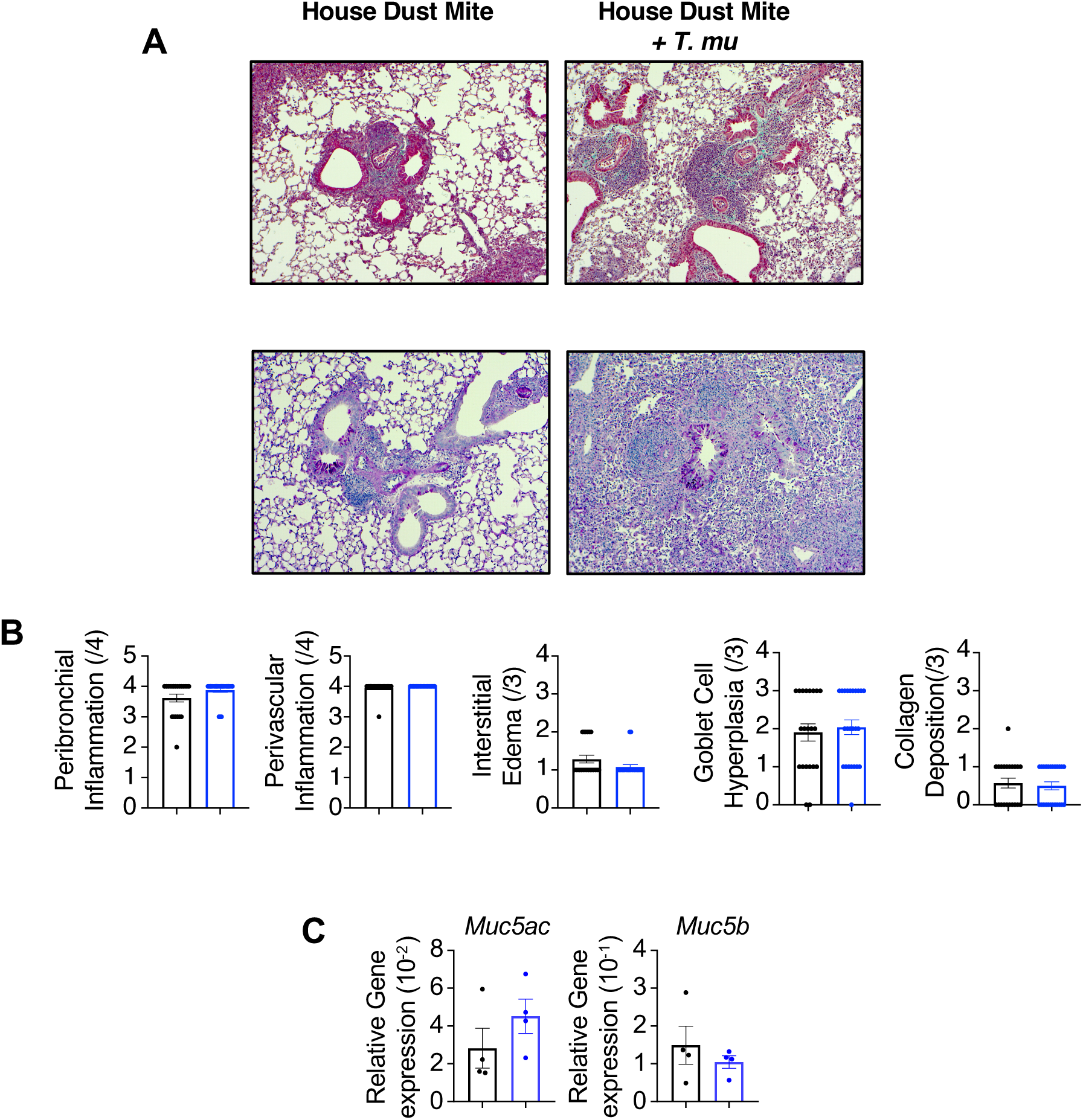
Exacerbated airway inflammation following colonization with *Tritrichomonas musculis*. (A) Masson’s trichrome (top) and PAS (bottom) stained lung sections of control and *T.mu* colonized mice following HDM-induced allergic airway inflammation. (B) Peribronchial and perivascular inflammation, interstitial edema, goblet cell hyperplasia and collagen deposition scores from lung histological analysis. (C) Lung mRNA expression of *Muc5ac* and *Muc5b* relative to *Gapdh.* Student’s *t* test was performed; statistical significance is indicated by *P<0.05.

## Graphical abstract

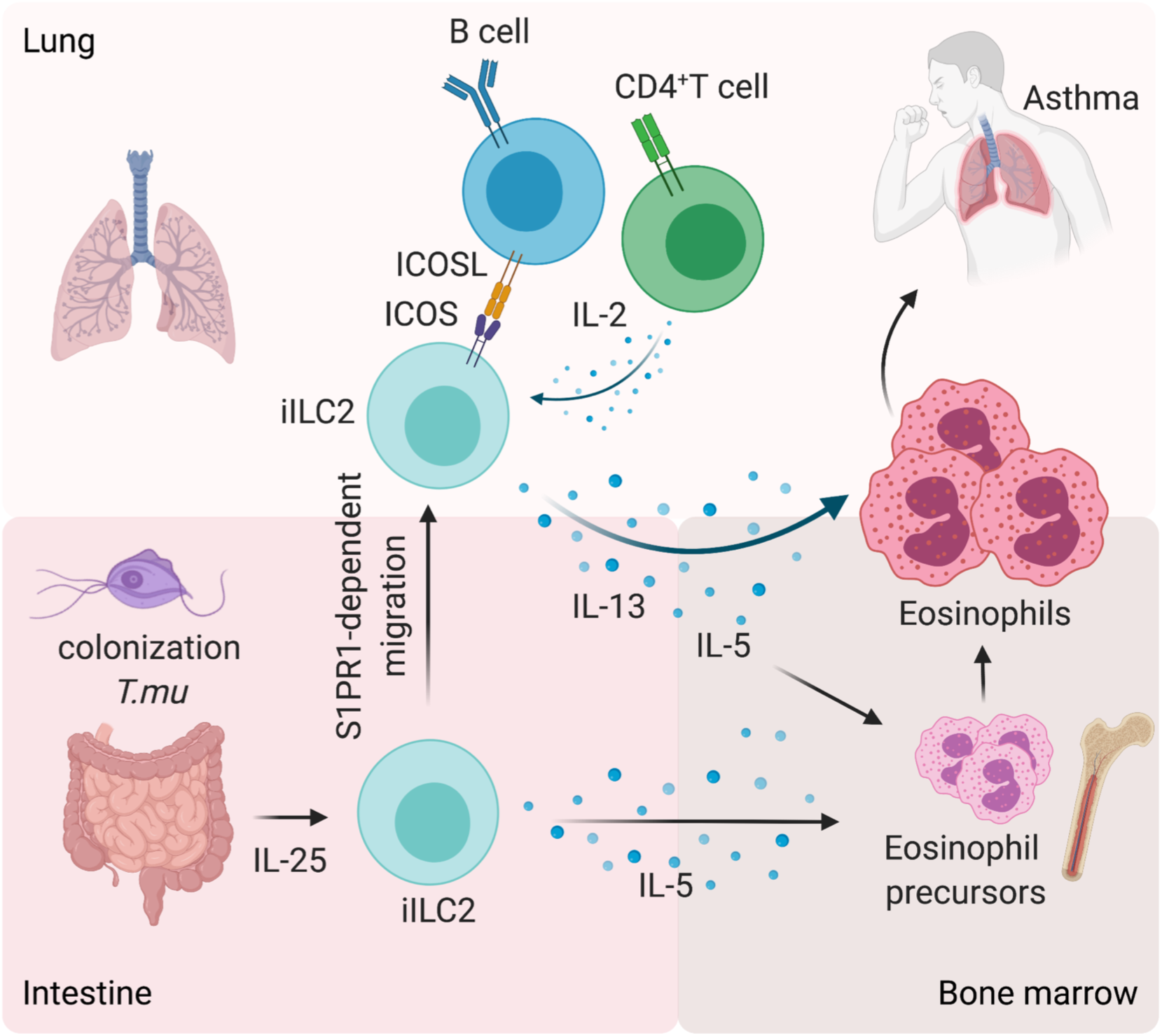

## Notes

### Competing Interest Statement

The authors have declared no competing interest.

## References

1. F. Sommer, F. Backhed, The gut microbiota--masters of host development and physiology. Nat Rev Microbiol 11, 227–238 (2013).

2. Q. Hamid, M. Tulic, Immunobiology of asthma. Annu Rev Physiol 71, 489–507 (2009).

3. M. C. Arrieta et al., Early infancy microbial and metabolic alterations affect risk of childhood asthma. Sci Transl Med 7, 307ra152 (2015).

4. W. Barcik, R. C. T. Boutin, M. Sokolowska, B. B. Finlay, The Role of Lung and Gut Microbiota in the Pathology of Asthma. Immunity 52, 241–255 (2020).

5. K. F. Budden et al., Emerging pathogenic links between microbiota and the gut-lung axis. Nat Rev Microbiol 15, 55–63 (2017).

6. A. T. Dang, B. J. Marsland, Microbes, metabolites, and the gut-lung axis. Mucosal Immunol 12, 843–850 (2019).

7. K. J. Blake, X. R. Jiang, I. M. Chiu, Neuronal Regulation of Immunity in the Skin and Lungs. Trends Neurosci 42, 537–551 (2019).

8. Y. Huang et al., S1P-dependent interorgan trafficking of group 2 innate lymphoid cells supports host defense. Science 359, 114–119 (2018).

9. K. Burrows, L. Ngai, F. Wong, D. Won, A. Mortha, ILC2 Activation by Protozoan Commensal Microbes. Int J Mol Sci 20, (2019).

10. C. Schneider, C. E. O’Leary, R. M. Locksley, Regulation of immune responses by tuft cells. Nat Rev Immunol 19, 584–593 (2019).

11. T. E. Billipp, M. S. Nadjsombati, J. von Moltke, Tuning tuft cells: new ligands and effector functions reveal tissue-specific function. Curr Opin Immunol 68, 98–106 (2021).

12. A. Chudnovskiy et al., Host-Protozoan Interactions Protect from Mucosal Infections through Activation of the Inflammasome. Cell 167, 444–456 e414 (2016).

13. M. R. Howitt et al., Tuft cells, taste-chemosensory cells, orchestrate parasite type 2 immunity in the gut. Science 351, 1329–1333 (2016).

14. M. S. Nadjsombati et al., Detection of Succinate by Intestinal Tuft Cells Triggers a Type 2 Innate Immune Circuit. Immunity 49, 33–41 e37 (2018).

15. Dientamoeba fragilis: A harmless commensal or a mild pathogen? Paediatr Child Health 3, 81–82 (1998).

16. P. Chiaranunt et al., NLRP1B and NLRP3 Control the Host Response following Colonization with the Commensal Protist Tritrichomonas musculis. J Immunol, (2022).

17. A. N. McKenzie, Type-2 innate lymphoid cells in asthma and allergy. Ann Am Thorac Soc 11 Suppl 5, S263–270 (2014).

18. A. Hamann, D. P. Andrew, D. Jablonski-Westrich, B. Holzmann, E. C. Butcher, Role of alpha 4-integrins in lymphocyte homing to mucosal tissues in vivo. J Immunol 152, 3282–3293 (1994).

19. A. A. Humbles et al., The murine CCR3 receptor regulates both the role of eosinophils and mast cells in allergen-induced airway inflammation and hyperresponsiveness. Proc Natl Acad Sci U S A 99, 1479–1484 (2002).

20. S. M. Pope, N. Zimmermann, K. F. Stringer, M. L. Karow, M. E. Rothenberg, The eotaxin chemokines and CCR3 are fundamental regulators of allergen-induced pulmonary eosinophilia. J Immunol 175, 5341–5350 (2005).

21. Y. Huang et al., IL-25-responsive, lineage-negative KLRG1(hi) cells are multipotential ’inflammatory’ type 2 innate lymphoid cells. Nat Immunol 16, 161–169 (2015).

22. D. Paclik, C. Stehle, A. Lahmann, A. Hutloff, C. Romagnani, ICOS regulates the pool of group 2 innate lymphoid cells under homeostatic and inflammatory conditions in mice. Eur J Immunol 45, 2766–2772 (2015).

23. T. Y. F. Halim et al., Tissue-Restricted Adaptive Type 2 Immunity Is Orchestrated by Expression of the Costimulatory Molecule OX40L on Group 2 Innate Lymphoid Cells. Immunity 48, 1195–1207 e1196 (2018).

24. F. Van Gool et al., Interleukin-5-producing group 2 innate lymphoid cells control eosinophilia induced by interleukin-2 therapy. Blood 124, 3572–3576 (2014).

25. C. J. Oliphant et al., MHCII-mediated dialog between group 2 innate lymphoid cells and CD4(+) T cells potentiates type 2 immunity and promotes parasitic helminth expulsion. Immunity 41, 283–295 (2014).

26. H. C. van Woerden et al., Association between protozoa in sputum and asthma: a case-control study. Respir Med 105, 877–884 (2011).

27. M. Mac Aogain et al., Metagenomics Reveals a Core Macrolide Resistome Related to Microbiota in Chronic Respiratory Disease. Am J Respir Crit Care Med 202, 433–447 (2020).

28. G. A. Mensah, J. P. Kiley, G. H. Gibbons, Generating evidence to inform an update of asthma clinical practice guidelines: Perspectives from the National Heart, Lung, and Blood Institute. J Allergy Clin Immunol 142, 744–748 (2018).

29. P. G. Woodruff et al., T-helper type 2-driven inflammation defines major subphenotypes of asthma. Am J Respir Crit Care Med 180, 388–395 (2009).

## References

1. K. Burrows, P. Chiaranunt, L. Ngai, A. Mortha, Rapid isolation of mouse ILCs from murine intestinal tissues. Methods Enzymol 631, 305–327 (2020).

2. M. Mac Aogain et al., Metagenomics Reveals a Core Macrolide Resistome Related to Microbiota in Chronic Respiratory Disease. Am J Respir Crit Care Med 202, 433–447 (2020).

